# Cows visually discriminate and cross-modally recognise familiar and unfamiliar human faces in videos

**DOI:** 10.1101/2025.07.20.665795

**Authors:** Océane Amichaud, Julie Lemarchand, Fabien Cornilleau, Plotine Jardat, Vitor H. B. Ferreira, Ludovic Calandreau, Léa Lansade

## Abstract

Social recognition has been studied and demonstrated in many species. In domesticated species, the long evolutionary history shared with humans has led to investigations into their cognitive abilities towards humans, particularly regarding discrimination and recognition of humans. The present study investigated whether cows are capable of visual discrimination and cross-modal recognition of familiar and unfamiliar humans. Thirty-two cows were exposed to two tests: a visual preference test, during which two silent videos were shown simultaneously - each displaying either a familiar or an unfamiliar human face - and a cross-modal test, during which the videos were accompanied by either a congruent or incongruent voice. During the visual preference test, cows looked significantly longer at the video showing the unfamiliar person (p = 0.025). In the cross-modal test, they looked significantly longer at the video that was congruent with the voice being played (p = 0.021). These two results show that cows are able to discriminate between familiar and unfamiliar individuals and form cross-modal representations of these people. Based on these results, future research should explore whether cows can adjust their behaviour depending on the person they are interacting with - a capacity that may reflect their agency in human–animal relationships.

## Introduction

Social recognition has been demonstrated in many animal species and plays a crucial role in regulating social interactions [1,2]. One of its fundamental prerequisites is the ability to discriminate between individuals [3], which supports the formation and maintenance of social bonds within groups. To achieve this, some species rely primarily on a dominant sensory modality, while others integrate multiple sensory channels such as visual, olfactory, and auditory cues [4].

Among these, the visual modality has been widely investigated in domestic species using two-dimensional facial images as stimuli. Facial perception is considered a key component of social recognition, as faces convey important information such as age, sex, and individual identity [5,6]. Several studies have shown that animals can discriminate their own species from other species [7–10], and can also distinguish between familiar and unfamiliar conspecifics based solely on facial cues [5,11–13]. To test the visual modality, several protocols are available, including choice-based tests relying on conditioning, as well as the preferential looking paradigm. The latter method involves comparing the gaze duration directed towards various stimuli, based on the premise that individuals will focus their visual attention longer on stimuli that are more relevant to them [14]. In a visual preference test, by choosing one stimulus over another, an individual demonstrates that they perceive differences between these stimuli, thereby indicating discrimination [15]. This test has shown, for instance, that when dogs are presented with pictures of familiar and unfamiliar dog faces, they display a preference for the familiar face by looking at it more, suggesting that they are able to discriminate familiar from unfamiliar pictures of conspecifics [13]. Animals can also discriminate conspecifics using only auditory or olfactory cues. For instance, dogs and horses can respectively distinguish conspecifics on the basis of their barking or urine samples [16,17].

All of these studies are based on the discrimination of conspecifics using a single sensory modality. Some studies have gone further, investigating whether animals are capable of associating current sensory cues with information previously acquired through different modalities [18]. This is known as cross-modal recognition. For example, Proops *et al*. [19] evaluated the ability of horses to individually recognise herd members cross-modally. To do this, they presented horses with a familiar conspecific and then played a vocalisation that had been recorded either from the horse that had just been seen (congruent trial) or from another horse (incongruent trial). Horses’ reaction times and gaze durations differed according to the type of trial, suggesting that horses possess a multimodal representation of familiar individuals.

Domestic species have a long evolutionary history with humans, and recent studies have focused on their interspecific socio-cognitive abilities towards us. In particular, researchers have notably examined the ability of domestic mammals to visually discriminate and recognise humans [20]. For instance, goats and chicks can distinguish familiar and unfamiliar human faces [21,22]. It has been shown that horses can discriminate fraternal and identical twins [23]. Sheep can recognise familiar and unfamiliar human faces [24]. Dogs recognise faces in photographs of their owners and horses can recognise a photograph of their keeper even if they have not seen them in six months, and even if the faces have been altered by changing their colour, covering their eyes, or modifying their hair [25,26]. Dogs and horses are also capable of cross-modally recognising men and women [27,28]. Horses and cats can also form cross-modal representations of humans [29–31].

Despite a growing body of evidence suggesting that domestic animals can recognise human faces, this ability has not yet been demonstrated in cows. At least one study has investigated this question, but concluded that cows were unable to discriminate between real humans when only their faces were visible in a conditioning-based discrimination test [32]. Given the number of species in which human recognition has been demonstrated, it would be surprising if this were not the case with cattle, thus warranting further investigation. Indeed, cattle are a good model for studying facial discrimination and recognition of familiar and unfamiliar humans. They are social animals and were domesticated 10,500 years ago [33]. They possess good visual acuity and a large visual field (330°) [34]. Moreover, previous studies have shown that cows can discriminate between individual people using multiple visual cues such as body height, colour of clothes and faces (under certain conditions) [32,35,36]. They have also been shown to recognise conspecifics in photographs [37].

Using two methodologies, progressing from the simplest to the most complex, the present study aims to test cows’ ability to: 1) visually discriminate between familiar and unfamiliar human faces (visual preference tests) and 2) cross-modally recognise familiar humans, by associating their voice with their face (cross-modal tests). For the visual preference tests, the hypothesis was that cows’ looking durations would differ between familiar and unfamiliar individuals. For the cross-modal tests, our hypothesis was that the animals would be able to associate the faces of familiar individuals with their voices. This would likely result in varying gaze durations depending on the congruence between the observed face and the heard voice. At this stage, it was difficult to predict the direction of these variations, since this is one of the first studies to include cows in such a paradigm, and the literature suggests that the direction of the effect can vary both within and between species. For instance, horses generally look longer at faces that are incongruent with the voice [30,31,38,39], while dogs generally do the opposite [40,41]. We also hypothesised that cows would experience different emotions in response to familiar and unfamiliar stimuli, which could result in differences in heart rate variation in response to the voices in the cross-modal tests.

## Materials and methods

### Ethics statement

This experiment was approved by the Val de Loire Ethical Committee (CEEA VdL, Nouzilly, France, authorisation number CE19 – 2025-2502-1). All the humans filmed in this study provided written authorisation for the use of their image.

### Subjects and housing

The study involved 34 Prim’ Holstein cows (*Bos taurus taurus*) aged 21.4 ± 15.2 months (mean ± sd) reared at the Experimental Unit PAO (UEPAO, 37,380 Nouzilly, France, https://doi. org/10. 15454/1. 55738 96321 728955E12), INRAE. These cows were housed in groups in indoor stalls enriched with automatic brushes and bedded with straw. Water was available ad libitum. They had been handled daily for feeding and care by four caretakers since birth, but they may also occasionally have encounter other individuals, such as students or colleagues visiting the farm.

### Experimental setup

The experiment took place in a barn housing cows not included in the study, adjacent to the home barn where the participating cows were kept. Before each test session, the experimenters ensured that no external sounds (human voices or farm machinery) were audible from the barn. The tested cow was placed in a test pen (l.159 x w.90 x h.192 cm; Fig 1). The videos were projected on two projection screens (l.216 x w.129 cm; dimensions of the projected image: l.110 x w.65cm) placed 280cm in front of the cow. Each test was filmed by three cameras (FDR-AX43A, Sony, Japan), one placed between the screens and the other two on both sides of the screens. The cows were fitted with a heart rate sensor belt (Polar Equine, Polar, Finland). The sensor was linked via Bluetooth to the Polar Beat smartphone application to display and record heart rate in real time.

**Fig 1.**
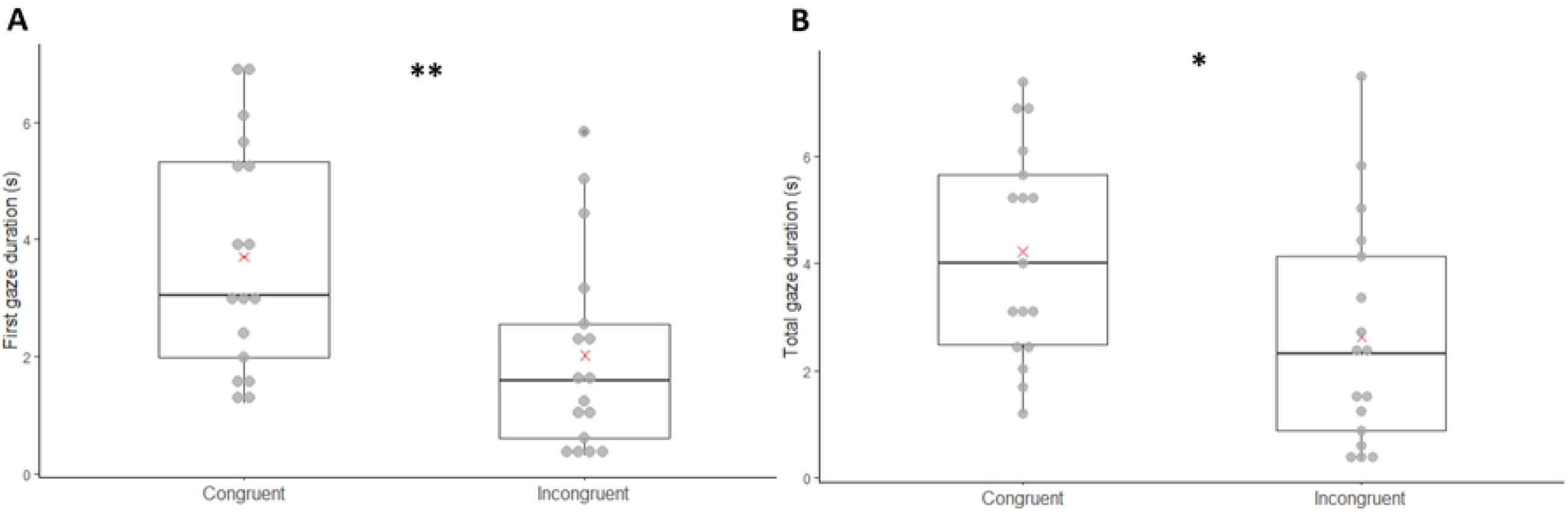
Experimental setup.

### Video preparation

During test sessions, two portrait videos were shown simultaneously (Fig 1): a video of a familiar man (a caretaker) and a video of an unfamiliar man (whom the cows had never seen before). Each video was shown on one of the screens. We selected only men to avoid discrimination on the basis of sex. Four familiar and four unfamiliar men were filmed. They were recorded under similar conditions, in the same room, at the same location within the room, with artificial light only and with the same framing (centred on the image and shoulders visible). All the men were recorded saying the following sentences with a neutral expression: « La réunion commence bientôt, je vais chercher mes affaires. Le chat monte dans l’arbre. », meaning « The meeting starts soon, I’m going to get my things. The cat is climbing the tree. ». These sentences were chosen because they contain words that cows are not used to hearing. Using Audacity software (v. 3.7.1, https://www.audacityteam.org/), the average intensity with which the men spoke was calculated from the audio of the recordings and the sound of each video was adjusted to this average value.

### Familiarisation

A familiarisation session, conducted prior to testing days, was used to familiarise the cows with the test environment. They were familiarised with crossing the outdoor courtyard to reach the testing stall. In the test pen, they were also familiarised with being fitted with the heart rate monitor belt. Once the cow was in position and equipped, she had to maintain a heart rate below 95 beats per minute (bpm) for two consecutive minutes to meet the familiarisation criterion. This threshold of 95 bpm was chosen because it is higher than the basal heart rate of young and adult bovids and has been exceeded following acute stress in several studies [42,43]. If this criterion was met, or if it was not achieved within five minutes, the cow was returned to her home pen (27 out of 34 cows met the criterion). A second familiarisation session, identical to the first, was conducted for each cow immediately prior to testing. If the cow maintained a heart rate below 95 bpm for two consecutive minutes, the test phase began immediately (28 cows). If the threshold was not reached within five minutes, the test was postponed to the following day, with another familiarisation session carried out beforehand (6 cows). Of the remaining six cows, four succeeded on their second attempt. The final two cows did not meet the criterion even after the second session and were excluded from testing, as the procedure was considered too stressful for them. In total, 32 cows completed the test phase, and analyses were conducted on these 32 individuals.

### Tests

Once the familiarisation criterion was met, the animals, equipped with the heart rate sensor belt underwent two successive tests: the visual preference test, immediately followed by the cross-modal test. This sequence was repeated a second time with different stimuli (faces and voice) after a 4-second interval during which black screens were projected.

### Visual preference test

The visual preference test lasted 8 seconds. Two muted videos, one showing a familiar face and the other an unfamiliar face, were presented simultaneously (Fig 1). Familiar and unfamiliar faces were randomly selected from a pool of four familiar and four unfamiliar individuals. The presentation side of the familiar and unfamiliar faces was also randomly assigned and varied between individuals and repetitions. We checked that this factor did not influence our results (see Statistical Analysis and S1 Table).

### Cross-modal test

Immediately after the visual preference test, the cows were subjected to a cross-modal recognition test for another 8 seconds. In addition to the same two videos presented during the visual preference test, an audio recording of a voice corresponding to either the familiar or the unfamiliar individual was broadcast by a loudspeaker (Megaboom 3, Ultimate Ears, United States; Fig 1) placed between the two screens and facing the cow. The sound was slightly offset from the image to prevent the cow from associating the voice she heard with the mouth movements of one of the people shown in the video. These voices were broadcast with an approximate intensity of 70 dB from where the cows’ head was, because it corresponded approximately to the intensity perceived when real people are talking. The presentation side of the congruent and incongruent videos, as well as the familiarity of the voice, were also randomly assigned and varied between individuals and repetitions. We also verified that this latter factor did not influence our results (see Statistical Analysis and S1 Table).

### Behavioural and physiological analyses

Videos of the tests were analysed with BORIS software [44] by the same coder. The screens were not visible on the cameras so that the coder did not know which side the familiar person and the congruent video appeared on. The time spent looking at each screen (right or left from the animal’s point of view) was quantified. The cow was considered to be looking at the right screen when her left eye was not fully visible to the right camera; conversely, she was considered to be looking at the left screen when her right eye was not fully visible to the left camera (Fig 1). For each cow, depending on the test and on the repetition, we obtained the total time spent watching the familiar and unfamiliar person and the time spent watching the congruent and incongruent video. First gaze duration was also measured for each stimulus (familiar/unfamiliar and congruent/incongruent). Both variables are commonly considered in similar tests [19,31,38], notably because significant effects can be observed only on one of these two variables [19].

Heart rate (HR) data were extracted via Polar Flow. These data were used 1) to monitor the cows’ emotional state during the familiarisation period and 2) to determine whether cows physiologically reacted to the voice they heard during cross-modal tests. For each cow, we removed HR values that were below 40 and above 180 bpm, as these values were considered artefactual [45]. For the cross-modal test of each repetition, mean HR and variation in HR (difference between the last 3 and first 3 seconds) were calculated for each cow.

### Statistical analyses

All statistical analyses were performed using R 4.4.2 [46] and all the figures presented in the results section were produced using the package *ggplot2* [47]. The significance threshold was fixed at p ≤ 0.05.

For the variables total gaze duration and first gaze duration, we excluded cows that looked at only one of the two screens in both repetitions, as such cases cannot be used to assess a visual preference. After applying the exclusion criteria, data from 22 cows were retained for the visual preference analysis, and from 17 cows for the cross-modal analysis, out of an initial sample of 32 cows. Moreover, for each cow we averaged the values over the two repetitions according to the familiarity of the person and according to the congruence of the video (depending on the test analysed). To determine whether the cows preferred a person and whether they were sensitive to congruence between voices and faces, the total gaze duration and first gaze duration for each screen were analysed according to the person presented (for the visual preference tests) and according to congruence (for the cross-modal tests). For this analysis we used generalised mixed effects models (GLMMs) from the *glmmTMB* package [48] with gaussian distributions. For each of the two response variables, two models were constructed (one per test): one to analyse the effect of the familiarity of the person presented and the other to analyse the effect of the congruence of the video. The identity of the cow was added as a random effect to account for individual variations in the paired data. Distributions, homoscedasticity of the residuals and the homogeneity of the variances were verified for the model fitting with the *DHARMa* package [49]. Additionally, the influence of the side presentation and the familiarity of voices (only for cross-modal tests) were checked, revealing that they did not influence the results (S1 Table).

To determine whether cows reacted to the voice they heard, the HR variation (difference between the last 3 and first 3 seconds) and the mean HR were analysed according to the voice broadcast during cross-modal tests (familiar vs unfamiliar voice). For this analysis, we used GLMMs from the *glmmTMB* package [48] with gaussian distributions. The identity of the cow was added as a random effect to account for individual variations in the paired data. Distributions, homoscedasticity of the residuals and homogeneity of the variances were verified for the model fitting with the *DHARMa* package [49].

## Results

### Visual preference tests

In the visual preference tests, the first gaze duration and the total gaze duration were significantly longer towards the unfamiliar person (GLMMs; respectively: χ^2^ = 5.120; df = 1; Z = 2.263; p = 0.024; Fig 2A; χ^2^ = 4.996; df = 1; Z = 2.235; p = 0.025; Fig 2B).

**Fig 2.**
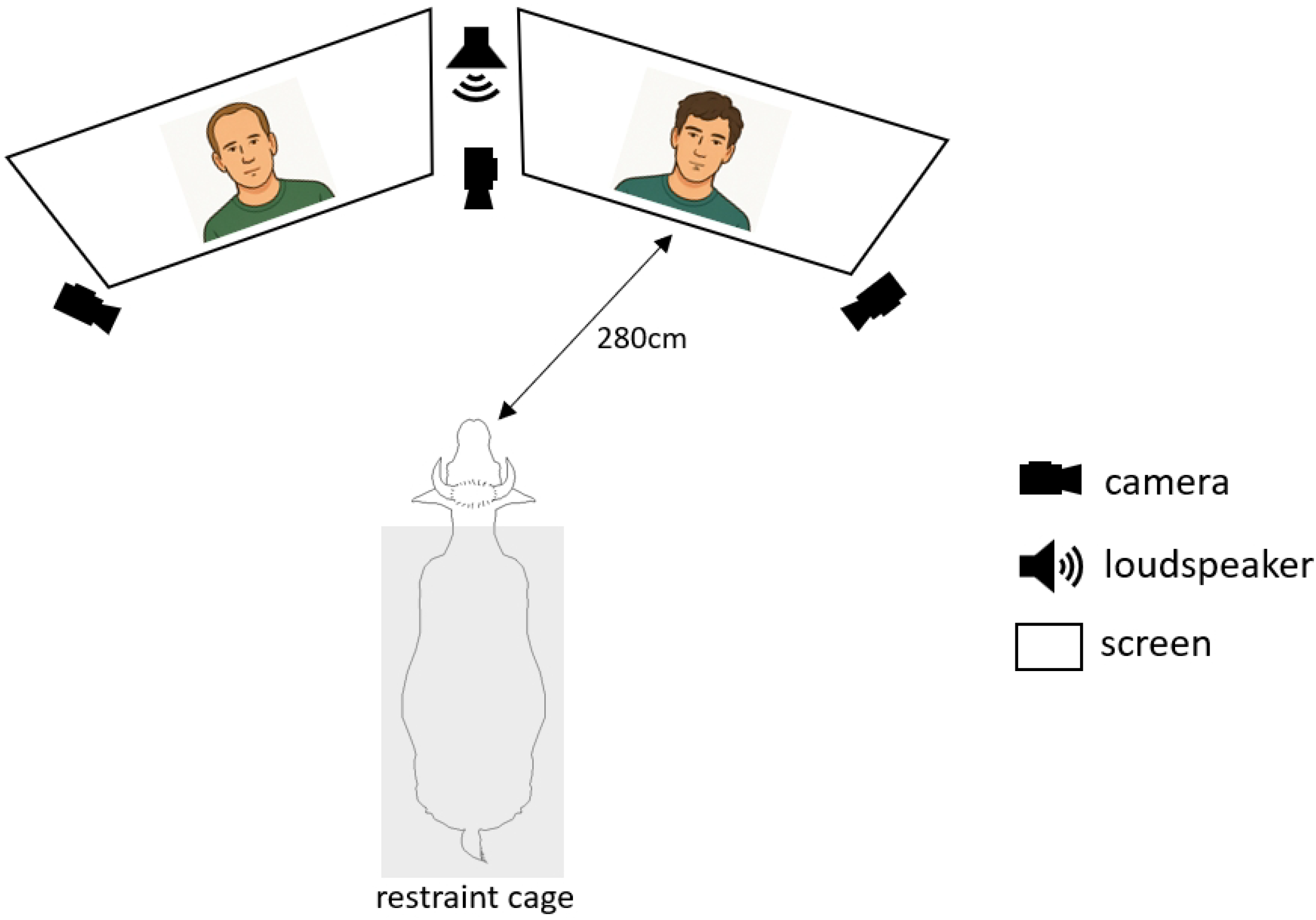
First gaze (A) and total gaze (B) duration according to the familiarity of the human face. Boxplots represent the median (central line), and the first and third quartiles (box limits). Red crosses represent averages. *: p ≤ 0.05, GLMMs, n=22.

### Cross-modal tests

First gaze duration and total gaze duration were significantly longer towards the congruent video (GLMMs; respectively: χ^2^ = 7.559; df = 1; Z = −2.749; p = 0.006; Fig 3A; χ^2^ = 5.351; df = 1; Z = −2.313; p = 0.021; Fig 3B).

**Fig 3.**
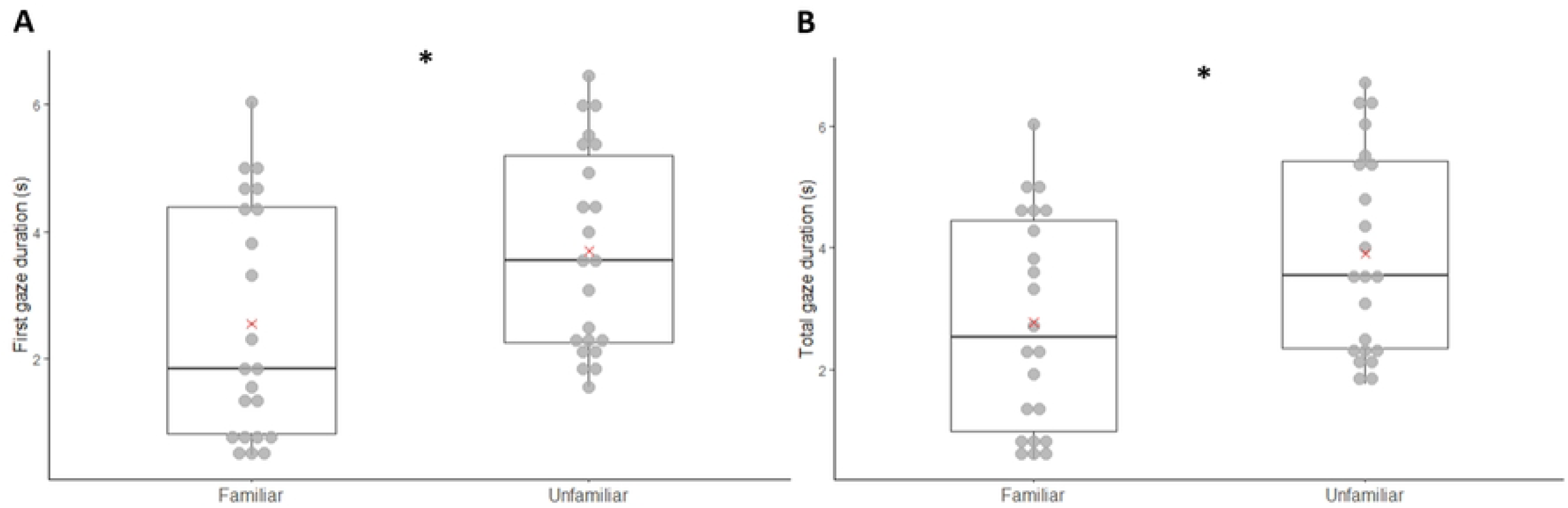
First gaze (A) and total gaze (B) duration according to the congruence of the video. Boxplots represent the median (central line), and the first and third quartiles (box limits). Red crosses represent averages. *: p ≤ 0.05, **: p ≤ 0.01, GLMMs, n=17.

The mean HR did not differ according to the voice broadcast (familiar voice: 84.84 ± 9.51 bpm; unfamiliar voice: 87.59 ± 11.51 bpm; GLMM; χ^2^ = 1.020; df = 1; Z = 1.010; p = 0.313). The variation in HR between the first 3 and last 3 seconds did not differ either (familiar voice: 1.02 ± 3.72 bpm; unfamiliar voice: 0.38 ± 5.08 bpm; GLMM; χ^2^ = 0.276; df = 1; Z = −0.525; p = 0.599).

## Discussion

The present study investigated whether cows are capable of visual discrimination and cross-modal recognition of familiar and unfamiliar humans. During the visual preference tests, cows looked significantly longer at the unfamiliar person, suggesting that they are able to discriminate between familiar and unfamiliar individuals using only a video of their faces as a cue. During the cross-modal tests, cows looked significantly longer at the face that matched the voice, indicating that they are able to associate familiar and unfamiliar voices with the corresponding face. However, they did not seem to show any physiological difference according to the voice heard, as the HR variables did not differ according to the familiarity of the voice.

### Visual preference tests

Using visual preference tests, we observed a difference between the first gaze duration and the total gaze duration directed at the unfamiliar person and the familiar person. More time spent watching a video indicates a preference for a given stimulus and suggests that the animal discriminates between the stimuli presented [50]. The observed results support the view that cows can categorise human faces according to familiarity. Thus, the capacity for differentiating human faces based on visual cues alone found in other domestic species [13,25] could extend to cows. Moreover, our results show that cows spent more time looking at the unfamiliar human, both in terms of first gaze duration and total gaze duration. Paradoxically, when exposed to photos of conspecifics’ heads, heifers spent more time looking at photos of familiar heifers than at photos of unfamiliar heifers [5]. The preference for looking at familiar or unfamiliar individuals appears to depend on whether the subject is a conspecific or a human. This is consistent with what was found in dogs: using a visual preference paradigm, Racca *et al*. [13] showed that dogs preferred to look at unfamiliar faces when presented with human faces, but preferred familiar faces when presented with conspecific faces. Unfamiliar human faces, due to their novelty, may be perceived as potentially threatening, thereby eliciting heightened vigilance and increased attentional allocation towards the stimulus.

What is particularly noteworthy in this test is that the human faces were presented only as two-dimensional videos, an artificial format that lacks depth cues and may alter colour perception. Moreover, no additional information such as scent, voice, body shape, posture, or gait was available to the animals. This indicates that cows are capable of recognising a person’s familiarity based solely on facial features.

### Cross-modal tests

Our results indicate that the recognition abilities of familiar humans seem to go beyond unimodal processing. In line with our hypothesis, cows watched the videos for varying lengths of time, depending on their congruence with the voice they heard. Cows’ ability to differentiate between familiar and unfamiliar humans in the videos, based on their congruence with the voice heard simultaneously, suggests that cows are capable of multimodal processing of human signals beyond facial or visual cues. More specifically, cows watched the congruent video for longer, i.e. the video presenting the person whose voice was being broadcast, for both first and total gaze durations. This longer gaze towards the congruent stimulus is in line with other studies using a cross-modal paradigm. For instance, Proops and McComb [4] tested whether horses were capable of individual recognition of familiar human handlers and showed that horses spent more time looking at the congruent stimulus. In contrast, studies investigating horses’ cross-modal representation of children and adults, or of human facial emotional expressions, as well as dogs’ and cats’ capacities to form cross-modal representation of individual humans, have shown that these animals spend more time looking (overall or for first gaze) at the stimulus that was incongruent with the broadcast sound [29–31,38,39,51]. These differences may be explained by variations in the species studied or in the experimental conditions, such as differences in the emotions or stress levels elicited by the presented stimuli.

Contrary to what we had predicted, the cows did not show any variation in heart rate in response to the voices. Several hypotheses can explain this result. Firstly, it is possible that cows discriminate between familiar and unfamiliar voices, but this does not necessarily imply that such discrimination is accompanied by a significant difference in emotional responses, particularly in terms of arousal level. It has been shown that goats are able to discriminate the emotional valence of human voices, however, this behavioural response was not accompanied by a significant physiological change [52]. Secondly, heart rate analyses were conducted during cross-modal tests, which involved relatively short durations of voice emission (eight seconds). This period may be too short to show any variation in heart rate. For instance, in a cross-modal test, Jardat *et al*. [30] showed that the horses’ heart rate increased when they heard a child’s voice, but their repetitions lasted twice as long (16 seconds). Moreover, in our study, the voice was not presented in isolation but simultaneously with two visual stimuli, which may have further reduced the likelihood of eliciting a clear physiological response to the sound. Thirdly, although the cows tested had never seen or heard the unfamiliar individuals presented during the test, they had all previously encountered new people (both visually and auditorily), including unfamiliar male voices. This prior exposure to novel social stimuli may have reduced their physiological reactivity to unfamiliar voices and could explain the absence of an increased heart rate response during the test. In such a context, where the stimulus does not elicit a strong emotional reaction, cows may be more inclined to direct their attention towards the congruent video, as observed in the present study.

This finding suggests that cows are capable of processing human cues and that they do not perceive all humans as a single, undifferentiated category, but are instead capable of distinguishing and recognising individuals they have previously met. Our results also indicate that cows are capable of integrating multiple sensory cues, reflecting a higher level of cognitive processing than that required for unimodal recognition, for example. Indeed, the ability to combine information from different sensory modalities suggests that cows form multisensory representations of individual humans. This study adds to a growing body of research showing that domesticated mammals develop complex socio-cognitive abilities in their interactions with humans [20]. Humans are part of their environment, particularly by providing them with daily care (e.g. by feeding or petting them) and animal welfare depends directly on the way in which an animal is able to perceive, interpret and analyse its environment. The recent definition of positive welfare emphasises the importance of experiencing mainly positive mental states, notably through the opportunity to make choices [53]. Indeed, for animals, the possibility of making choices can have a direct positive impact on their emotional state by giving them a feeling of control and agency [54]. In the case of the human-animal relationship, this is illustrated, for example, by giving the animal a choice as to when and how to interact [55]. In this way, the socio-cognitive abilities mentioned above can have an adaptive value and enable animals to adjust their behaviour according to the person’s profile.

## Conclusions

In this study, using visual preference and cross-modal tests, we showed that cows are able to process human faces presented in 2D on videos and to associate familiar and unfamiliar faces with the corresponding voices by integrating multiple sensory modalities. Such cognitive abilities highlights the complexity of human perception in domestic animals, as discussed in the review by Jardat and Lansade [20]. In cows, this innovative experimental design provides a promising tool for investigating a wider range of cognitive abilities in this species, notably their ability to recognise individual humans and to develop preferential interactions. A better understanding of how cows perceive and differentiate humans could help inform husbandry practices that incorporate human–animal interactions aligned with their cognitive abilities, in order to provide them with greater opportunities for choice and initiative in their relationship with humans - thereby reinforcing their agency, a key component of positive welfare [53,56].

## Acknowledgments

The authors would like to thank Ludovic Metivier, Eric Briant, Mickaël Dupont and David Georget from the UEPAO (Unité Expérimentale de Physiologie Animal de l’Orfrasière, https://doi. org/10. 15454/1. 55738 96321 728955E12) for the daily care of the animals during the experiment, for their technical help and for agreeing to be filmed for the videos we presented. We would also like to thank the four men who agreed to be filmed playing the role of unfamiliar people.

## Supporting information

**S1 Table. Evaluation of the influence of the presentation side of the familiar stimuli and the familiarity of voices (only for cross-modal tests) during the experiment**. Additional versions of the model were added to the analysis of visual preference and cross-modal tests. They included the variable presentation side of the familiar stimuli in interaction with the familiarity of the person (for visual preference analysis) and with the congruence (for cross-modal analysis). For cross-modal analysis only, they also included the familiarity of voices in interaction with the congruence of the video. Bold text highlights the models selected by the ANOVA because they significantly differed from the null model and/or significantly minimized the AIC value (difference > 2). The variables presentation side of familiar stimuli and voice familiarity were never included in the selected models, showing that they did not influence the results. The selected models corresponded to the ones previously built and selected (see Statistical analysis), therefore, the initial analysis was retained and is presented in the Results section.

## Notes

### Competing Interest Statement

The authors have declared no competing interest.

## References

1. Colgan PW. Comparative social recognition. New York: Wiley; 1983.

2. Ligout S, Porter RH. La reconnaissance sociale chez les mammifères : mécanismes et bases sensorielles impliquées. INRAE Prod Anim. 2006;19: 119–134. doi:10.20870/productions-animales.2006.19.2.3490

3. Tibbetts EA, Dale J. Individual recognition: it is good to be different. Trends Ecol Evol. 2007;22: 529–537. doi:10.1016/j.tree.2007.09.001

4. Proops L, McComb K. Cross-modal individual recognition in domestic horses (Equus caballus) extends to familiar humans. Proc R Soc B Biol Sci. 2012;279: 3131–3138. doi:10.1098/rspb.2012.0626

5. Coulon M, Baudoin C, Heyman Y, Deputte BL. Cattle discriminate between familiar and unfamiliar conspecifics by using only head visual cues. Anim Cogn. 2011;14: 279–290. doi:10.1007/s10071-010-0361-6

6. Behrmann M, Avidan G. Face perception: computational insights from phylogeny. Trends Cogn Sci. 2022;26: 350–363. doi:10.1016/j.tics.2022.01.006

7. Autier-Dérian D, Deputte BL, Chalvet-Monfray K, Coulon M, Mounier L. Visual discrimination of species in dogs (Canis familiaris). Anim Cogn. 2013;16: 637–651. doi:10.1007/s10071-013-0600-8

8. Coulon M, Deputte BL, Heyman Y, Delatouche L, Richard C, Baudoin C. Visual discrimination by heifers (Bos taurus) of their own species. J Comp Psychol. 2007;121: 198–204. doi:10.1037/0735-7036.121.2.198

9. Kendrick KM, Atkins K, Hinton MR, Broad KD, Fabre-Nys C, Keverne B. Facial and vocal discrimination in sheep. Anim Behav. 1995;49: 1665–1676. doi:10.1016/0003-3472(95)90088-8

10. Ragonese G, Baragli P, Mariti C, Gazzano A, Lanatà A, Ferlazzo A, et al. Interspecific two-dimensional visual discrimination of faces in horses (Equus caballus). PLOS ONE. 2021;16: e0247310. doi:10.1371/journal.pone.0247310

11. Kendrick KM, Atkins K, Hinton MR, Heavens P, Keverne B. Are faces special for sheep? Evidence from facial and object discrimination learning tests showing effects of inversion and social familiarity. Behav Processes. 1996;38: 19–35. doi:10.1016/0376-6357(96)00006-X

12. Langbein J, Moreno-Zambrano M, Siebert K. How do goats “read” 2D-images of familiar and unfamiliar conspecifics? Front Psychol. 2023;14. doi:10.3389/fpsyg.2023.1089566

13. Racca A, Amadei E, Ligout S, Guo K, Meints K, Mills D. Discrimination of human and dog faces and inversion responses in domestic dogs (Canis familiaris). Anim Cogn. 2010;13: 525–533. doi:10.1007/s10071-009-0303-3

14. Wilson VAD, Bethell EJ, Nawroth C. The use of gaze to study cognition: limitations, solutions, and applications to animal welfare. Front Psychol. 2023;14. doi:10.3389/fpsyg.2023.1147278

15. Koba R, Izumi A. Japanese monkeys (Macaca fuscata) discriminate between pictures of conspecific males and females without specific training. Behav Processes. 2008;79: 70–73. doi:10.1016/j.beproc.2008.04.005

16. Molnár C, Pongrácz P, Faragó T, Dóka A, Miklósi Á. Dogs discriminate between barks: The effect of context and identity of the caller. Behav Processes. 2009;82: 198–201. doi:10.1016/j.beproc.2009.06.011

17. Hothersall B, Harris P, Sörtoft L, Nicol CJ. Discrimination between conspecific odour samples in the horse (Equus caballus). Appl Anim Behav Sci. 2010;126: 37–44. doi:10.1016/j.applanim.2010.05.002

18. Pitcher BJ, Briefer EF, Baciadonna L, McElligott AG. Cross-modal recognition of familiar conspecifics in goats. R Soc Open Sci. 2017;4: 160346. doi:10.1098/rsos.160346

19. Proops L, McComb K, Reby D. Cross-modal individual recognition in domestic horses (Equus caballus). Proc Natl Acad Sci. 2009;106: 947–951. doi:10.1073/pnas.0809127105

20. Jardat P, Lansade L. Cognition and the human–animal relationship: a review of the sociocognitive skills of domestic mammals toward humans. Anim Cogn. 2022;25: 369–384. doi:10.1007/s10071-021-01557-6

21. Deutsch J, Lebing S, Eggert A, Nawroth C. Goats who stare at video screens – assessing behavioural responses of goats towards images of familiar and unfamiliar con- and heterospecifics. Peer Community J. 2024;4. doi:10.24072/pcjournal.473

22. Wood SMW, Wood JN. Face recognition in newly hatched chicks at the onset of vision. J Exp Psychol Anim Learn Cogn. 2015;41: 206–215. doi:10.1037/xan0000059

23. Stone SM. Human facial discrimination in horses: can they tell us apart? Anim Cogn. 2010;13: 51–61. doi:10.1007/s10071-009-0244-x

24. Knolle F, Goncalves RP, Morton AJ. Sheep recognize familiar and unfamiliar human faces from two-dimensional images. R Soc Open Sci. 2017;4: 171228. doi:10.1098/rsos.171228

25. Lansade L, Colson V, Parias C, Trösch M, Reigner F, Calandreau L. Female horses spontaneously identify a photograph of their keeper, last seen six months previously. Sci Rep. 2020;10: 6302. doi:10.1038/s41598-020-62940-w

26. Eatherington CJ, Mongillo P, Lõoke M, Marinelli L. Dogs (Canis familiaris) recognise our faces in photographs: implications for existing and future research. Anim Cogn. 2020;23: 711–719. doi:10.1007/s10071-020-01382-3

27. Gouyet C, Ringhofer M, Yamamoto S, Jardat P, Parias C, Reigner F, et al. Horses cross-modally recognize women and men. Sci Rep. 2023;13: 3864. doi:10.1038/s41598-023-30830-6

28. Ratcliffe VF, McComb K, Reby D. Cross-modal discrimination of human gender by domestic dogs. Anim Behav. 2014;91: 127–135. doi:10.1016/j.anbehav.2014.03.009

29. Takagi S, Arahori M, Chijiiwa H, Saito A, Kuroshima H, Fujita K. Cats match voice and face: cross-modal representation of humans in cats (Felis catus). Anim Cogn. 2019;22: 901–906. doi:10.1007/s10071-019-01265-2

30. Jardat P, Ringhofer M, Yamamoto S, Gouyet C, Degrande R, Parias C, et al. Horses form cross-modal representations of adults and children. Anim Cogn. 2023;26: 369–377. doi:10.1007/s10071-022-01667-9

31. Lampe JF, Andre J. Cross-modal recognition of human individuals in domestic horses (Equus caballus). Anim Cogn. 2012;15: 623–630. doi:10.1007/s10071-012-0490-1

32. Rybarczyk P, Koba Y, Rushen J, Tanida H, de Passillé AM. Can cows discriminate people by their faces? Appl Anim Behav Sci. 2001;74: 175–189. doi:10.1016/S0168-1591(01)00162-9

33. Bollongino R, Burger J, Powell A, Mashkour M, Vigne J-D, Thomas MG. Modern Taurine Cattle Descended from Small Number of Near-Eastern Founders. Mol Biol Evol. 2012;29: 2101–2104. doi:10.1093/molbev/mss092

34. Adamczyk K, Górecka-Bruzda A, Nowicki J, Gumułka M, Edyta M, Schwarz T, et al. Perception of environment in farm animals – A review. Ann Anim Sci. 2015;15: 565–589. doi:10.1515/aoas-2015-0031

35. Taylor AA, Davis H. Individual humans as discriminative stimuli for cattle (Bos taurus). Appl Anim Behav Sci. 1998;58: 13–21. doi:10.1016/S0168-1591(97)00061-0

36. Munksgaard L, De Passillé AM, Rushen J, Thodberg K, Jensen MB. Discrimination of People by Dairy Cows Based on Handling1. J Dairy Sci. 1997;80: 1106–1112. doi:10.3168/jds.S0022-0302(97)76036-3

37. Coulon M, Deputte BL, Heyman Y, Baudoin C. Individual Recognition in Domestic Cattle (Bos taurus): Evidence from 2D-Images of Heads from Different Breeds. PLOS ONE. 2009;4: e4441. doi:10.1371/journal.pone.0004441

38. Jardat P, Liehrmann O, Reigner F, Parias C, Calandreau L, Lansade L. Horses discriminate between human facial and vocal expressions of sadness and joy. Anim Cogn. 2023;26: 1733–1742. doi:10.1007/s10071-023-01817-7

39. Trösch M, Cuzol F, Parias C, Calandreau L, Nowak R, Lansade L. Horses Categorize Human Emotions Cross-Modally Based on Facial Expression and Non-Verbal Vocalizations. Animals. 2019;9: 862. doi:10.3390/ani9110862

40. Albuquerque N, Guo K, Wilkinson A, Savalli C, Otta E, Mills D. Dogs recognize dog and human emotions. Biol Lett. 2016;12: 20150883. doi:10.1098/rsbl.2015.0883

41. Yong MH, Ruffman T. Domestic dogs match human male voices to faces, but not for females. Behaviour. 2015;152: 1585–1600. doi:10.1163/1568539X-00003294

42. Hopster H, O’Connell JM, Blokhuis HJ. Acute effects of cow-calf separation on heart rate, plasma cortisol and behaviour in multiparous dairy cows. Appl Anim Behav Sci. 1995;44: 1–8. doi:10.1016/0168-1591(95)00581-C

43. Van Reenen CG, O’Connell NE, Van der Werf JTN, Korte SM, Hopster H, Jones RB, et al. Responses of calves to acute stress: Individual consistency and relations between behavioral and physiological measures. Physiol Behav. 2005;85: 557–570. doi:10.1016/j.physbeh.2005.06.015

44. Friard O, Gamba M. BORIS: a free, versatile open-source event-logging software for video/audio coding and live observations. Methods Ecol Evol. 2016;7: 1325–1330. doi:10.1111/2041-210X.12584

45. Laister S, Stockinger B, Regner A-M, Zenger K, Knierim U, Winckler C. Social licking in dairy cattle—Effects on heart rate in performers and receivers. Appl Anim Behav Sci. 2011;130: 81–90. doi:10.1016/j.applanim.2010.12.003

46. R Core Team. R: A language and environment for statistical computing. Vienna, Austria: R Foundation for Statistical Computing; 2024. Available: https://www.R-project.org/

47. Wickham H. Programming with ggplot2. In: Wickham H, editor. ggplot2: Elegant Graphics for Data Analysis. Cham: Springer International Publishing; 2016. pp. 241–253. doi:10.1007/978-3-319-24277-4_12

48. Brooks ME, Kristensen K, Benthem KJ van, Magnusson A, Berg CW, Nielsen A, et al. Modeling zero-inflated count data with glmmTMB. bioRxiv; 2017. p. 132753. doi:10.1101/132753

49. Hartig F. DHARMa: Residual Diagnostics for Hierarchical (Multi-Level /Mixed) Regression Models. 2024. Available: https://CRAN.R-project.org/package=DHARMa

50. Watanabe S, Shinozuka K, Kikusui T. Preference for and discrimination of videos of conspecific social behavior in mice. Anim Cogn. 2016;19: 523–531. doi:10.1007/s10071-016-0953-x

51. Adachi I, Kuwahata H, Fujita K. Dogs recall their owner’s face upon hearing the owner’s voice. Anim Cogn. 2007;10: 17–21. doi:10.1007/s10071-006-0025-8

52. Mason MA, Semple S, Marshall HH, McElligott AG. Goats discriminate emotional valence in the human voice. Anim Behav. 2024;209: 227–240. doi:10.1016/j.anbehav.2023.12.008

53. Rault J-L, Bateson M, Boissy A, Forkman B, Grinde B, Gygax L, et al. A consensus on the definition of positive animal welfare. Biol Lett. 2025;21: 20240382. doi:10.1098/rsbl.2024.0382

54. Englund MD, Cronin KA. Choice, control, and animal welfare: definitions and essential inquiries to advance animal welfare science. Front Vet Sci. 2023;10. doi:10.3389/fvets.2023.1250251

55. McGowan RTS, Bolte C, Barnett HR, Perez-Camargo G, Martin F. Can you spare 15 min? The measurable positive impact of a 15-min petting session on shelter dog well-being. Appl Anim Behav Sci. 2018;203: 42–54. doi:10.1016/j.applanim.2018.02.011

56. Špinka M. Animal agency, animal awareness and animal welfare. Anim Welf. 2019;28: 11–20. doi:10.7120/09627286.28.1.011

